# Genome-wide consensus transcriptional signatures identify synaptic pruning linking Alzheimer’s disease and epilepsy

**DOI:** 10.1101/2024.10.28.618752

**Authors:** Huihong Li, Zhiran Xie, Yuxuan Tian, Ruoyin Zhou, Yaxi Yang, Bingying Lin, Si Chen, Jie Wu, Zihan Deng, Jianwei Li, Mingjie Chen, Xueke Liu, Yushan Sun, Bing Huang, Naili Wei, Xiaoyu Ji

**Author notes:** **Corresponding authors** Contact: Dr. Xiaoyu Ji.

## Abstract

Alzheimer’s disease (AD) and epilepsy (EP) share a complex bidirectional relationship, yet the molecular mechanisms underlying their comorbidity remain insufficiently explored. To identify potential transcriptional programs across animal models and human patients with AD and EP, we conducted a comprehensive genome-wide transcriptomic analysis. Our investigation included mouse models of temporal lobe epilepsy (pilocarpine- and kainic acid-induced; n = 280), AD transgenic models (7 transgenic models expressing human tau or amyloid pathology; n = 257), and performed cross-species validation in human cohorts (EP: n = 182; AD: n = 301). We identified a highly conserved immune-related module across all models and patient cohorts. The hub consensus signatures of this module were centered around a microglial synaptic pruning pathway involving *TYROBP*, *TREM2*, and *C1Q* complement components. Gene regulatory network analysis identified *TYROBP* as the key upstream hub signature. These signatures showed consistent up-regulation in both conditions and strong diagnostic potential. Differential expression analyses revealed their predominant expression in specific microglial subpopulations associated with complement-mediated synaptic pruning and immune activation. Neural circuit modeling further demonstrates the asymmetric sensitivity of synaptic pruning to network dynamics. Loss of inhibitory synapses has a disproportionately significant impact on neural network excitation/inhibition balance and synchronization. Our findings support microglial complement-mediated synaptic pruning as a conserved central pathway linking neurodegeneration to epileptogenesis, suggesting a promising therapeutic target for AD and EP comorbidity.

**Graphic abstract:** 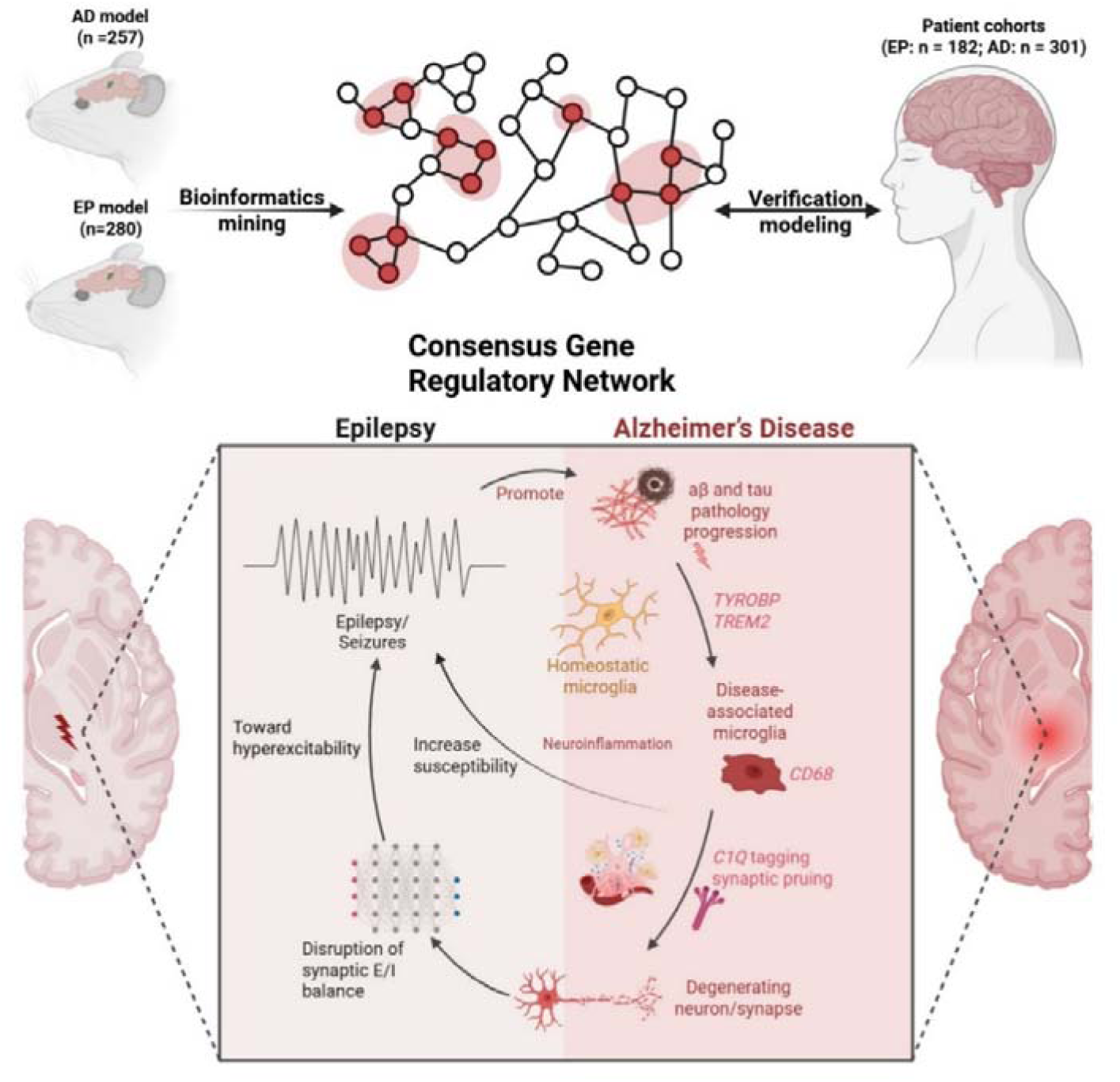

**Highlights:**

- Genome-wide transcriptomic analysis across epilepsy (EP) and Alzheimer’s disease (AD) models and patients identified a conserved immune module.
- The *TYROBP-TREM2-C1Q* microglial synaptic pruning pathway was identified as a central consensus signature across model and patient cohorts.
- Consensus signatures possess potential diagnostic utility in AD and EP patients, with predominant expression in specific microglial subpopulations
- Neural circuit modeling demonstrates the asymmetric effects of synaptic pruning, whereby loss of inhibitory synapses disproportionately disrupts the E/I balance, leading to increased network synchronization.

## 1. Introduction

Alzheimer’s disease (AD) and epilepsy (EP) are two common neurological diseases with a complex bidirectional relationship that contributes to their respective pathogenesis (1–3). AD, characterized by progressive cognitive decline and neurodegeneration, is the leading cause of dementia worldwide, affecting an estimated 55 million individuals. This number is projected to triple by 2050 as life expectancy increases (4). EP, characterized by recurrent unprovoked seizures, affects over 50 million individuals worldwide, with an exceptionally high incidence among elderly populations, especially those with neurodegenerative conditions such as AD (5,6).

Epidemiological studies demonstrate a strong association between AD and EP. AD patients exhibit increased seizure susceptibility, with 10–22% experiencing unprovoked seizures, while 22–54% demonstrate subclinical epileptiform activity during disease progression (7,8). These seizures, predominantly occurring in the temporal lobe, correlate with accelerated cognitive decline and exacerbated AD pathology (9). Conversely, EP patients demonstrate a two- to three-fold increased risk of developing dementia (10,11). This elevated risk is particularly evident in temporal lobe epilepsy (TLE), where numerous pathological features overlap with AD (9).

The pathological overlap between AD and EP manifests across multiple biological scales (12). Both conditions feature amyloid deposition, tau accumulation, and hippocampal sclerosis at the molecular and cellular levels, particularly in TLE (13–15). Chronic seizures increase neuronal activity and metabolic demands, potentially accelerating the production and aggregation of amyloid-β (Aβ) and hyperphosphorylated tau protein (16,17). Conversely, the progressive accumulation of these pathological proteins in AD appears to lower seizure thresholds, promoting hyperexcitability and epileptiform activity (18,19). At the network level, both disorders demonstrate profound disruption of the excitation/inhibition (E/I) balance, with a shift toward hyperexcitability driving both cognitive impairment and seizure susceptibility. This network dysfunction is accompanied by impaired synaptic plasticity, altered synchronization patterns, and progressive circuit reorganization (19–21).

Despite these established clinical and pathological connections, the precise molecular mechanisms governing the AD-EP relationship remain insufficiently explored. While previous studies have identified potential contributors—including dysregulated protein homeostasis, mitochondrial dysfunction, and chronic inflammatory responses—most investigations have examined these conditions in isolation rather than explicitly addressing their comorbidity (22–24). Several factors have limited current research approaches: 1) inadequate integration of transcriptomic data across multiple disease models and human cohorts; 2) insufficient attention to shared cellular processes that bridge molecular alterations with network dysfunction; and 3) limited application of computational modeling to connect molecular findings with clinically relevant physiological outcomes.

The challenge of treating EP in AD patients further highlights the need for deeper mechanistic insights. Clinicians face complex decisions regarding seizure treatment in AD patients, including whether to treat subclinical epileptiform activity and which antiseizure medications minimize cognitive side effects (25). Emerging therapeutic approaches require a more precise understanding of the shared pathological mechanisms. Systems biology and computational neuroscience offer powerful tools to integrate insights from animal models and human patients, potentially revealing actionable therapeutic targets that simultaneously address both conditions.

This study investigated the molecular mechanisms underlying the comorbidity between AD and EP through genome-wide transcriptional analyses. Our approach included: 1) comprehensive transcriptomic analysis of mouse models of TLE (n = 280) and AD (n = 257) to identify consensus signatures across disease paradigms, 2) cross-species validation in human cohorts (EP: n = 182; AD: n = 301) to assess clinical relevance, and 3) computational modeling to connect molecular findings with network-level mechanisms. Through these integrated analyses, we aimed to identify conserved transcriptional programs that might explain the mechanistic link between neurodegeneration and epileptogenesis, potentially revealing novel therapeutic targets for both conditions.

## 2. Materials and Methods

A workflow overview of the datasets and analyses can be found in the figure below.

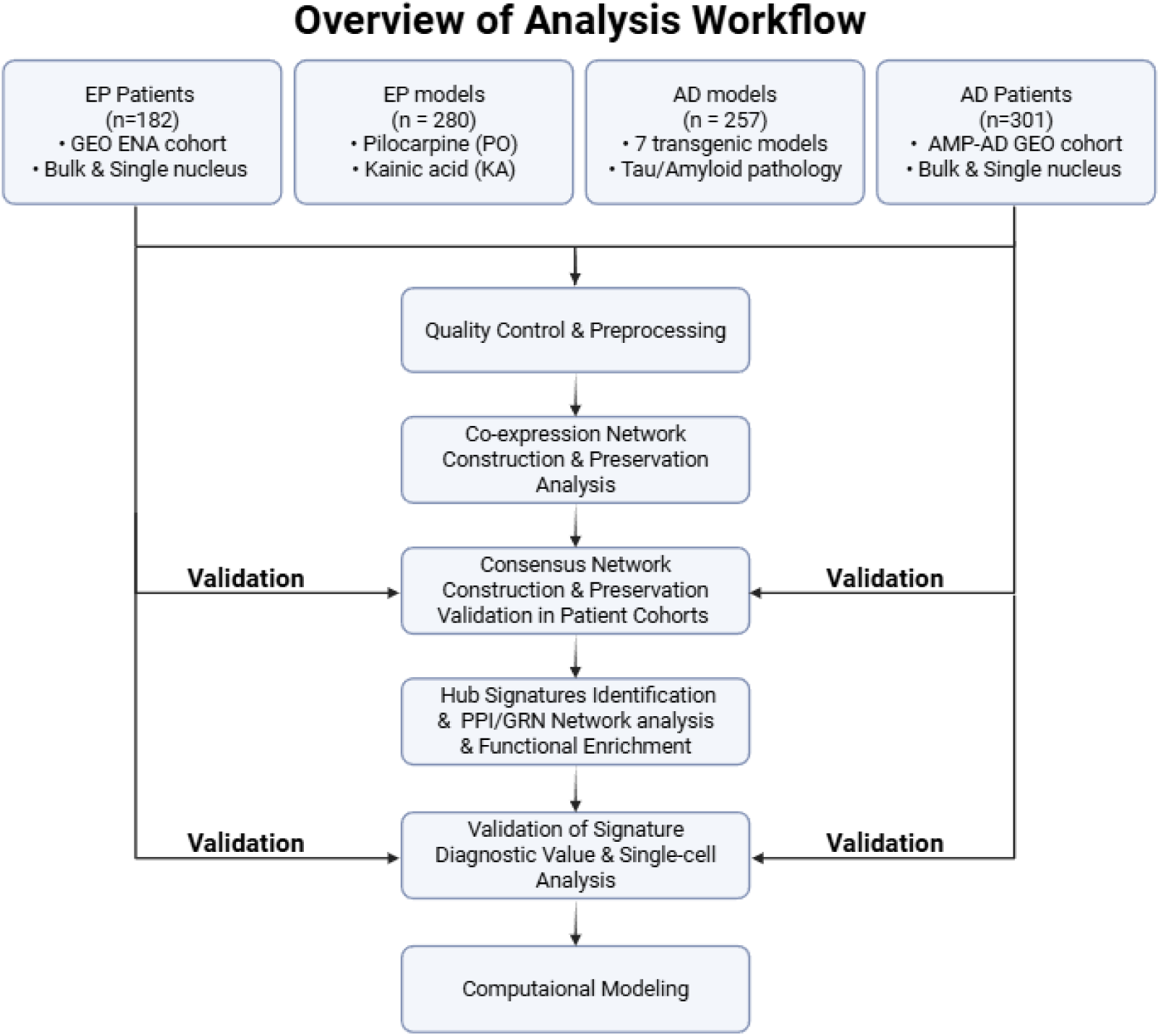

### 2.1 Data Collection

We retrieved several large-scale, well-characterized datasets from EP and AD research that focused on genome-wide expression patterns in well-characterized disease models and appropriate controls. The EP model datasets included two of the most used TLE modeling approaches. The first dataset (PRJEB18790) contained 200 hippocampi of pilocarpine-induced (PO) model mice and wild-type (WT) controls (26). The second dataset (GSE73878) comprised 80 hippocampal samples from the kainic acid-induced (KA) model and WT control. For the AD model datasets, we selected two datasets that systematically examined genome-wide expression alterations in amyloid and tau pathology progression. The Matarin dataset (GSE64398) contained 113 hippocampal samples from five mouse models and their controls: TAU mice, TAS10 mice, TPM mice, and TASTPM mice (both heterozygous and homozygous variants) (27). The Castanho dataset (GSE125957) included J20 and rTg4510 models with 144 hippocampal and entorhinal cortex samples and matched controls (28). To validate disease co-expression modules across species, we constructed modules using 129 hippocampal samples from TLE patients (GSE63808) and incorporated 30 consensus AD modules from the AMP-AD consortium’s meta-analysis of 2,114 samples across seven brain regions (29,30). To evaluate the diagnostic value of transcriptional signatures, we analyzed 47 hippocampal samples from TLE patients and controls (E-MTAB-3123) and 253 samples from AD patients and controls (GSE48350). In addition, we reanalyzed single-nucleus RNA-sequencing (snRNA-seq) datasets to characterize cell-type and subtype-specific expression of identified signatures. Details about data collection, quality control, preprocessing, parameters, and additional methods can be found in **Text S1**.

### 2.2 Gene Co-expression Networks Construction

To construct and compare gene co-expression networks (GCNs) between EP and AD model datasets, we implemented the standard meta-WGCNA comparative methodology (31,32), effectively identifying conserved gene modules while mitigating batch effects between datasets. Using the WGCNA package (1.73), we constructed signed GCNs based on genome-wide expression profiles. Network construction employed the minimum soft-thresholding power that achieved scale-free topology (*R²* ≥ 0.85) across all model datasets. Using the signed network option, we generated adjacency matrices and transformed them into topological overlap matrices (TOM) to quantify gene connectivity profile similarities. Module identification proceeded through average linkage hierarchical clustering of TOM dissimilarity (1-TOM), followed by dynamic tree cutting (parameters: deepSplit = 1, minClusterSize = 27). We also constructed consensus modules to identify GCNs consistently in all conditions. This consensus approach involved calculating the component-wise minimum values of the topological overlap matrices from all datasets. We calculated module eigengenes (MEs) from the first principal component to summarize module expression profiles for each module. We then identified disease-associated modules by computing Pearson correlation coefficients between MEs and genotype information, with significance thresholds of *P* < 0.05 and |correlation coefficient| > 0.35. Additionally, we determined module membership (MM) and genotype significance (GS) by calculating Pearson correlations between individual genes and their respective MEs or genotypes. Gene Ontology (GO) enrichment analyses for module annotation were conducted using clusterProfiler (4.10.1) (33). Hub genes act as key drivers within a module and were identified using stringent criteria to minimize false positives: genes with MM > 0.9 and ranking in the top 5% for intramodular connectivity within each module. Consensus hub signatures were obtained by intersecting the hub genes of key modules across different datasets.

### 2.3 Module Preservation Analysis

To evaluate the preservation of EP model modules across AD model datasets, we implemented dual analytical approaches as described by Langfelder et al. (34). First, we calculated Zsummary preservation statistics using the modulePreservation function. This composite metric comprehensively assesses preservation by integrating module size, density, and connectivity parameters, quantifying how effectively EP module structures are maintained in AD datasets. We interpreted the preservation evidence according to established thresholds: strong (Zsummary > 10), moderate (2 < Zsummary < 10), or absent (Zsummary < 2). Additionally, we performed cross-tabulation analysis to quantify module overlap between EP and AD model datasets, as well as between model and patient datasets. This approach generated contingency tables detailing the genes shared between the reference and test dataset modules. We determined overlap significance using Fisher’s exact test with Benjamini-Hochberg (BH) correction for multiple testing.

### 2.4 Protein-Protein Interaction and Gene Regulatory Network Analysis

To explore interactions and potential regulatory relationships among consensus hub signatures, we constructed a protein-protein interaction (PPI) network using the STRING database (12.0) with a confidence score threshold of 0.4 and a false discovery rate (FDR) of 5% (35). We also performed regulatory network analysis with the GENIE3 (1.28.0) algorithm, which applies a random forest approach to infer potential regulatory relationships from gene expression data. GENIE3 ranks regulatory interactions by weight, with higher weights indicating stronger regulatory links (36). Gene expression profiles of the consensus hub signatures from all datasets were used to infer plausible regulatory interactions. Only the top 30% plausible regulatory links were retained to construct gene regulatory networks to reduce false positives.

### 2.5 Single-nucleus RNA-seq Analysis

snRNA-seq datasets from EP and AD cohorts were reanalyzed to investigate cell-type-specific transcriptional changes of consensus transcriptional signatures. For the EP cohort, we utilized the snRNA-seq dataset (GSE190453) profiling temporal lobe tissue from TLE patients and non-epileptic controls (cells = 66305) (37). For the AD cohort (syn51758062.1), we analyzed snRNA-seq data from the Religious Orders Study and Rush Memory and Aging Project (ROSMAP) cohort, encompassing 48 prefrontal cortex (PFC) samples from AD patients and non-pathology controls (cells = 50985) (38). Data preprocessing was performed using the Seurat package (5.0.1) according to the original studies (39). Dimensionality reduction was conducted through PCA, with the first 30 components used for clustering and UMAP visualization. Cell-type annotation followed the definitions established in the original studies and calculated scores for each nucleus using the AddModuleScore() function, and cell-type identity was assigned based on the highest-scoring module. Differential expressions of consensus transcriptional signature between control and disease groups were evaluated using two-sided Wilcoxon rank-sum tests. To demonstrate the expression of our signatures in microglial subtypes, we utilized the comprehensive single-cell microglial dataset (GSE204702), which covers 215,680 microglia from different neurological diseases and brain regions (40). We performed differential expression analysis for their 12 subclasses and analyzed the expression patterns of our consensus signatures across the identified microglial subtypes.

### 2.6 Neural Circuit Modeling

We implemented a spiking neural network using the Izhikevich neuron model to simulate neural circuit dynamics and explore the contribution of synaptic pruning to pathological changes in AD and EP. The network consisted of 1000 standard Izhikevich neurons (800 excitatory, 200 inhibitory) and was simulated for 1000ms using the Brian2 package (41). Network architecture included random sparse connectivity (0.1 probability) with conductance-based synapses modeling AMPA (excitatory) and GABA-a (inhibitory) currents across four connection types: E→E, E→I, I→E, and I→I (42). Synaptic weights were drawn from a standard normal distribution and normalized to [0,1]. We simulated synaptic pruning by selectively removing excitatory or inhibitory synapses with the lowest weights (0-50%, with 5% increments), repeating each simulation at least 50 times. To quantify network dynamics, we calculated the excitation/inhibition (E/I) ratio and event synchrony (ES) (43). The E/I ratio compared total excitatory currents to the absolute magnitude of inhibitory currents, while ES measured spike timing coordination between excitatory neuron pairs within 5-ms windows. Both metrics were normalized to baseline unpruned values to reflect pruning-induced changes. Statistical analysis included Pearson correlation coefficients and linear regression to quantify excitatory and inhibitory pruning effect sizes on network dynamics. Complete details of the computational model parameters, equations, and analysis methods can be found in **Text S2.**

## 3. Results

### 3.1 Identification of Consensus Gene Co-Expression Network and Hub Signatures

To identify conserved molecular signatures underlying EP and AD, we employed an unbiased, genome-wide approach to construct GCNs across four independent model datasets: two EP models (pilocarpine-induced [EP-PO] and kainic acid-induced [EP-KA]) and two AD model datasets (AD-Castanho: [rTg4510, J20]; AD-Matarin: [TAU, TAS10, TPM, HO-TASTPM, HET-TASTPM]). After removing only genes with negligible or undetectable expressions, we retained all expressed genes (ranging from 12,558 to 17,813) to avoid selection bias. We identified common and distinct GCN modules across all model datasets (**Fig. 1A, S1**): EP-PO revealed 19 modules (97-1203 genes), EP-KA contained 14 modules (36-1534 genes), AD-Castanho yielded 28 modules (55-820 genes), and AD-Matarin showed 16 modules (33-1354 genes). Correlation analysis between module eigengenes and disease status identified disease-associated modules in each model (**Table S2**). In the EP-PO dataset, 12 modules were significantly associated with the EP genotype (|*r*| > 0.35, *P* < 0.05). Similarly, EP-KA had two disease-associated modules, AD-Castanho contained 12 modules correlated with tau pathology, with four modules correlated with general AD pathology, and AD-Matarin revealed five modules associated with tau pathology and two modules related to amyloid pathology, with three modules correlated with general AD pathology (**Fig. 1A, Table S2**).

**Figure 1.**
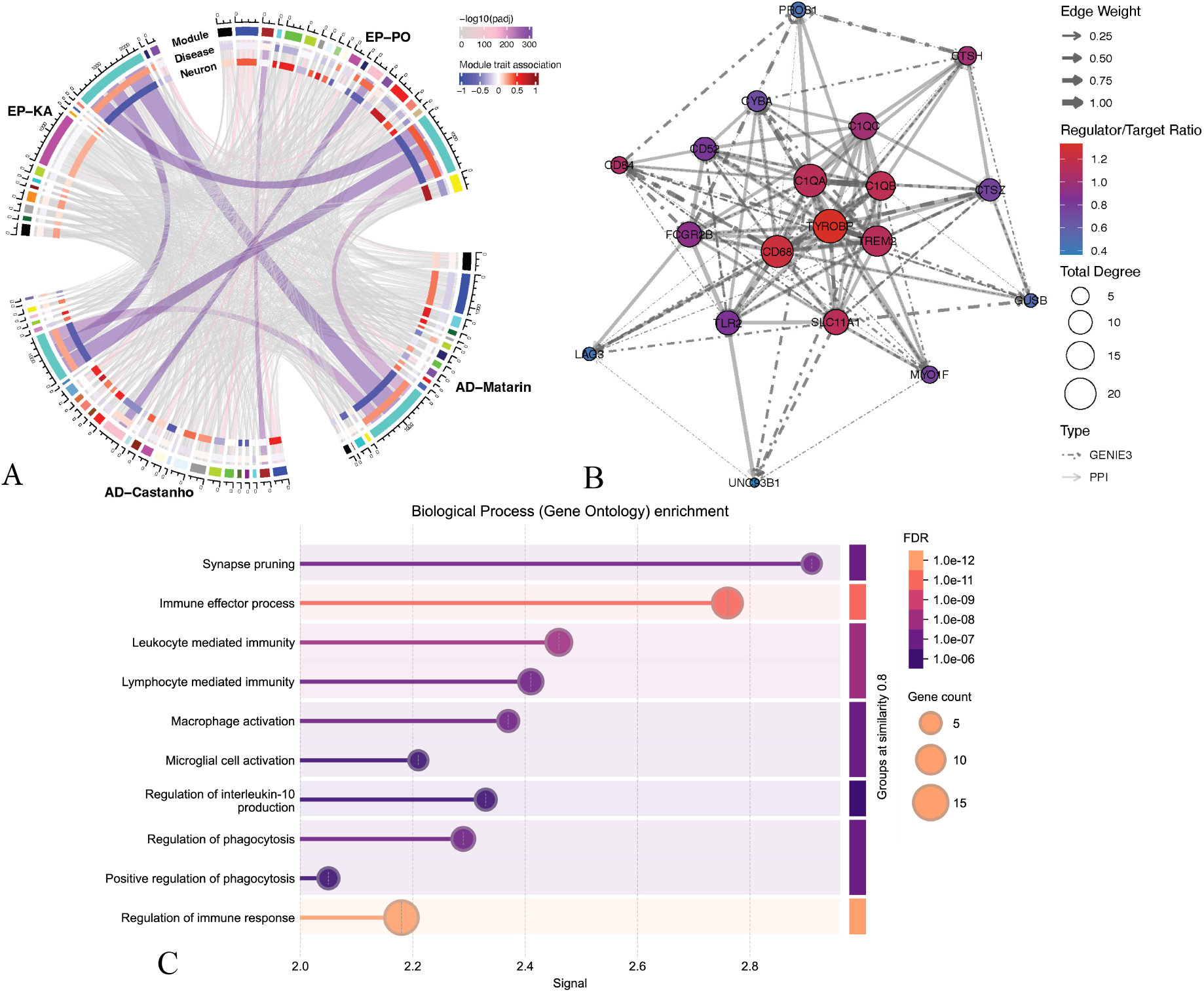
Identification of conserved immune-related GCNs and hub signatures. **(A)** Chord diagram showing module preservation across EP and AD models. Each segment represents a gene module identified in different datasets. Purple connections indicate statistically significant module overlap (Fisher’s exact test, adjusted *P <* 1E-300), with color intensity corresponding to significance level (-log10(padj)). Inner tracks display module correlations with disease status (red: positive correlation; blue: negative correlation) and neuron proportion. **(B)** Integrated PPI and regulatory network of consensus hub signatures identified across all models. Nodes represent hub genes consistently present in the preserved immune module, with node size proportional to total weighted degree. Solid lines indicate validated protein-protein interactions (STRING), dotted lines indicate computationally inferred gene regulation (GENIE3), with edge width reflecting interaction strength. Node color indicates regulator/target ratio, with red showing stronger regulatory potential. **(C)** GO enrichment analysis of consensus hub signatures. Bar length corresponds to enrichment signal strength, color indicates significance level (FDR), and size reflects gene count within each category. Bars represent significantly enriched biological processes, with “Synaptic pruning” emerging as the most enriched process with the highest signal.

Despite model and disease-specific differences, cross-condition preservation analysis revealed remarkable conservation of an immune module across all datasets (**Fig. 1A**). Using Z-summary preservation statistics and cross-tabulation approaches, we identified a large immune-related module (turquoise) with exceptionally highest preservation (mean Z-summary = 70 & mean median rank = 3 & Fisher’s adjusted *P* < 1E-300) across all pairwise comparisons (**Fig. 1A, Table S2**). This module was negatively correlated with deconvoluted neuron proportion and positively correlated with disease progression genotype in each dataset (|*r*| > 0.46, *P* < 9.31E-07) (**Table S2, Text S1**). The strong preservation pattern of this immune module across both diseases prompted us to construct consensus GCNs by integrating network topology information from all datasets (**Fig. S2**). We identified a robust immune-related and disease-associated consensus module containing 1617 genes common to all models with 19 consensus hub signatures (**Fig. 1B, Table S2-3**).

Among these 19 consensus hub signatures, PPI and gene regulatory network centrality analysis identified *TYROBP* as the most upstream hub signature with the highest total degree (regulator/target ratio = 1.33, total degree = 24.19), followed by *CD68*, *TREM2*, *C1Q* complement components, and *SLC11A1* (**Fig. 1B, Table S2**). GO enrichment analysis of the hub signature network identified “synaptic pruning” (*C1Q* complement components, *TREM2*) as the most significantly enriched biological process with the strongest signal (**Fig. 1C, Table S2**).

Taken together, our genome-wide transcriptomic analysis revealed a highly conserved immune module across EP and AD models, with a consensus regulatory network centered around *TYROBP*, *TREM2*, *CD68*, and *C1Q* components that coordinate microglia activation and synaptic pruning. This consensus molecular pathway represents a potential central mechanism linking both neurological conditions.

### 3.2 Consensus GCNs Significantly Overlap with GCNs in Human Patients

To validate the translational relevance of our identified consensus GCN, we compared it with disease-associated GCNs from patient cohorts. We employed a cross-species approach to identify overlaps between our consensus modules and AD-associated GCN modules from 2,114 samples across seven brain regions, as well as our newly constructed EP patient modules from 127 EP patient hippocampal samples (**Fig. 2A, S3, Table S3**).

**Figure 2.**
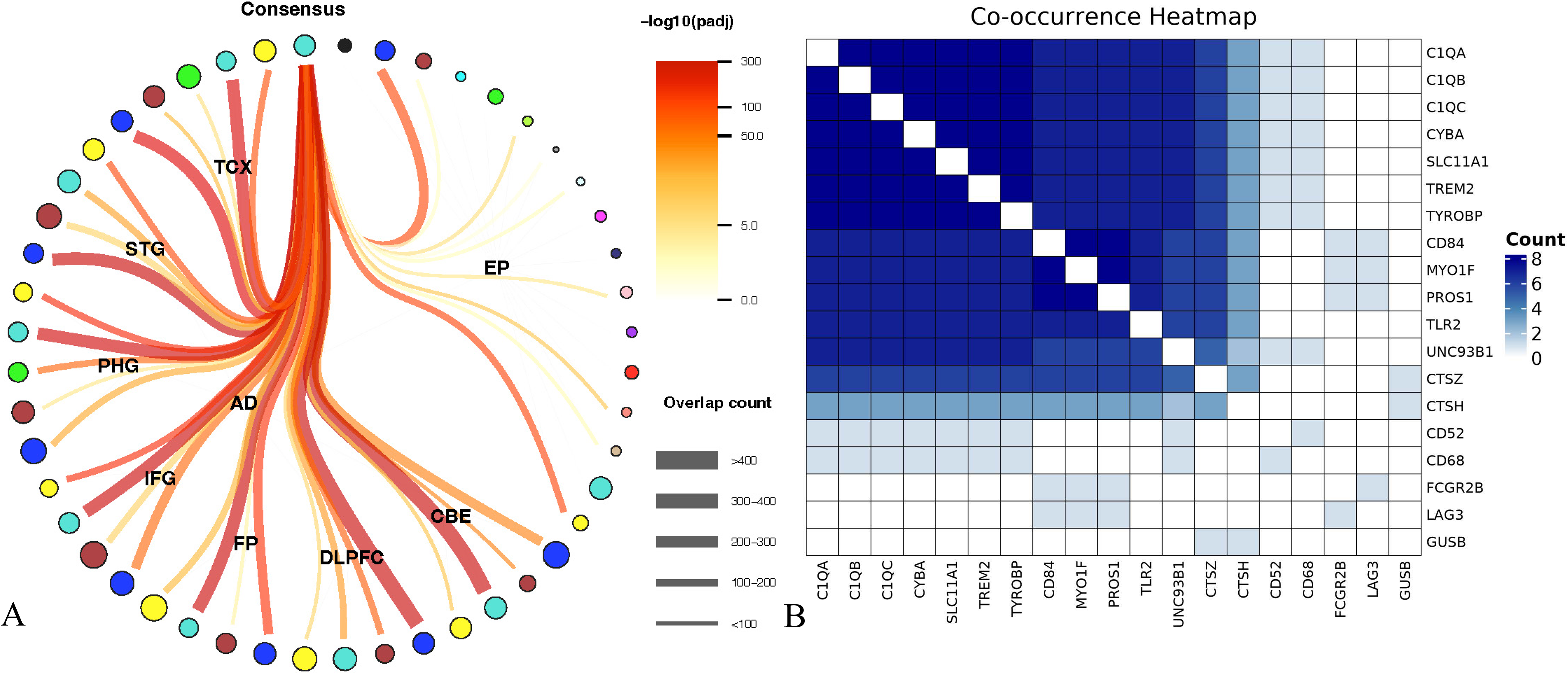
Comparison of consensus modules with patient modules. **(A)** The overlap between the consensus immune module and disease-associated GCNs from human patients. Peripheral nodes represent modules from different brain regions in AD and EP patient cohorts. Line thickness indicates the number of overlapping genes between each patient module and the consensus module, with thicker lines showing stronger connections. Line color intensity corresponds to statistical significance (-log10(adjusted *P*)), with red-orange hues indicating the highest significance levels. The significant associations (adjusted *P <* 0.05) were observed with modules across brain regions in AD patients, and with yellow and blue modules in EP patients. **(B)** Co-occurrence heatmap of the 19 identified consensus hub signatures across human disease modules. Color intensity represents the frequency of gene co-occurrence (darker blue indicates higher co-occurrence counts).

Our analysis revealed significant overlaps between the consensus immune module and multiple human AD modules across different brain regions. The strongest associations were observed with modules from the cerebellum (CBE), dorsolateral prefrontal cortex (DLPFC), and superior temporal gyrus (STG) regions. The consensus module demonstrated a powerful overlap with the “ CBE-turquoise” module (n = 502, 25.39% of genes, *P* < 1E-300), the “DLPFC-blue” module (n = 502, 28.67% of genes, *P* < 1E-300) and “STG-blue” (n = 389, 33.22% of genes, *P* < 1E-300) (**Fig. 2A, Table S2**). Similarly, substantial overlaps were detected with modules from the inferior frontal gyrus (IFG), superior temporal gyrus (STG), temporal cortex (TCX), and frontal pole (FP), with the overlap percentage varying from 2% to 33% (**Table S2**). For the EP patient cohort, our consensus module showed significant overlap with two EP patients’ GCN modules (yellow: n = 153, 39.64% of genes, *P* < 7.07E-88; blue: n = 279, 22.85% of genes, *P* < 1.06E-84) (**Fig. 2A, Table S2**). Notably, the top hub genes identified in our animal model analysis—*TYROBP*, *TREM2*, and the *C1Q* components—were also identified as central hub genes (top 2 intramodular connectivity) in the EP patient’s yellow module (**Table S3**). To further validate our hub consensus signatures findings, we examined whether the 19 previously identified hub signatures remained clustered within the same modules in human disease networks (**Fig. 2B**). The co-occurrence analysis revealed strong co-expression patterns among these signatures, with particularly strong co-occurrence observed among *C1Q* complement components, *TREM2*, *TYROBP*, *CYBA*, *and SLC11A1*. These genes showed the highest co-occurrence frequency (dark blue in the heatmap), suggesting they function as a coordinated unit across species and disease conditions.

Collectively, these cross-species preservation analyses demonstrate that the identified immune modules and hub signatures represent conserved biological entities relevant to both AD and EP pathogenesis in humans.

### 3.3 Consensus Hub Signatures Showed Microglial Specificity Upregulation

To validate the clinical relevance of our identified consensus hub signatures, we investigated their expression patterns in bulk and single-nucleus human cohorts and explored their distribution across microglial subtypes.

Expression profiling across patient cell types demonstrated that these signatures were predominantly localized to microglia (**Fig. 3A-D**), though some genes, including *CTSH*, *MYO1F*, and *GUSB*, exhibited broader expression across multiple cell types. In general, nearly all signatures showed significant upregulation in the EP cohort, while most were also upregulated in AD patient cohorts, with particularly elevated expression in microglia from late-stage pathology (**Fig. 3E-H**). Furthermore, diagnostic models developed by various machine learning algorithms based on bulk expression profiles of 19 consensus features achieved excellent discrimination between patients and controls (EP: AUC > 0.9, AD: AUC > 0.7). (**Fig. S4, Text S1**). Notably, *TYROBP*, *TREM2*, and *C1Q* complement genes—central consensus signatures involved in microglial synaptic pruning—demonstrated consistent upregulation in both disorders (**Fig. 3E-H**).

**Figure 3.**
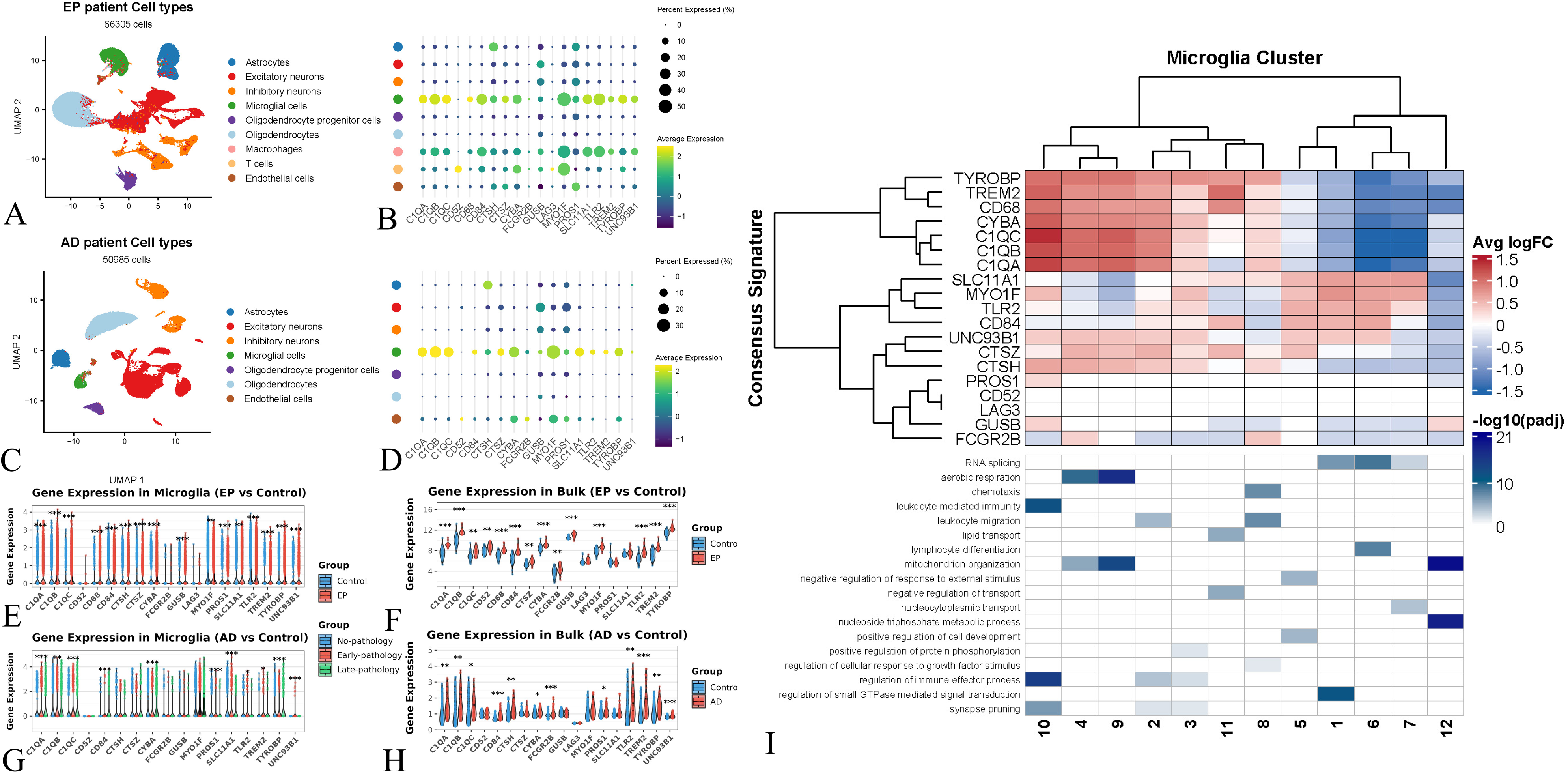
Expression of consensus hub signatures in EP and AD patient cohorts. **(A-D)** UMAP visualization and dot plot showing cell type-specific expression of consensus hub genes in EP patient samples **(A-B)** and AD patient samples **(C-D)**. Microglia show the highest expression of most hub genes, with *TYROBP*, *TREM2*, and *C1Q* complement genes exhibiting powerful expression. **(E-H)** Differential expression analysis of consensus hub genes in microglia and bulk tissue from EP patients (**E-F**) and AD patients (**G-H**) compared to controls. Bar heights represent fold changes with statistical significance indicated (*p < 0.05, **p < 0.01, ***p < 0.001). (**I**) Heatmap showing the expression patterns (average log fold change) of 19 consensus gene signatures across 12 distinct microglial clusters derived from Tuddenham’s data. The color scale represents average log fold change (logFC) or statistical significance for GO enrichment analysis. The dendrogram shows hierarchical clustering of both genes and microglial subtypes.

To understand the specific microglial contributions, we examined the expression patterns of our consensus signatures across microglial subtypes using a comprehensive single-cell RNA-sequencing dataset from Tuddenham et al., encompassing 215,680 microglial cells from 74 donors. Distinct expression patterns of consensus hub signatures across 12 defined microglial subtypes were found (**Fig. 3I**). Key signatures, including *TYROBP*, *TREM2*, *CD68*, and *C1Q* complement genes, showed prominent upregulation in clusters 10, 4, and 9, while being downregulated in clusters 12, 7, 6, and 1. According to GO functional characterization, these high-expressing clusters represent distinct microglial states: cluster 10 is enriched for genes involved in antigen presentation and complement-mediated synaptic pruning, while clusters 4 and 9 share genes associated with oxidative phosphorylation and immune response pathways (**Fig. 3I**). In contrast, clusters showing downregulation of our consensus signatures (clusters 1, 6, and 7) represent alternative microglial states characterized by metabolic and transcriptional regulatory functions.

These findings demonstrate that consensus hub signatures, particularly those involved in microglial activation and synaptic pruning, possess potential diagnostic utility and are expressed explicitly in functionally distinct microglial subpopulations. Immune-activated microglial subtypes, especially cluster 10, are primary mediators of synaptic pruning, potentially contributing to the comorbidity of AD and EP.

### 3.4 Asymmetric Effects of Synaptic Pruning on Neural Network Dynamics

To better understand how synaptic pruning influences neural network dynamics and its potential role in the comorbidity of AD and EP, we developed a spiking neural network model simulating the neural circuit. In this model, we systematically varied the extent of synaptic pruning (0% to 50%) at excitatory or inhibitory synapses and quantified changes in the E/I ratio and synchronization levels (**Fig. 4**).

**Figure 4.**
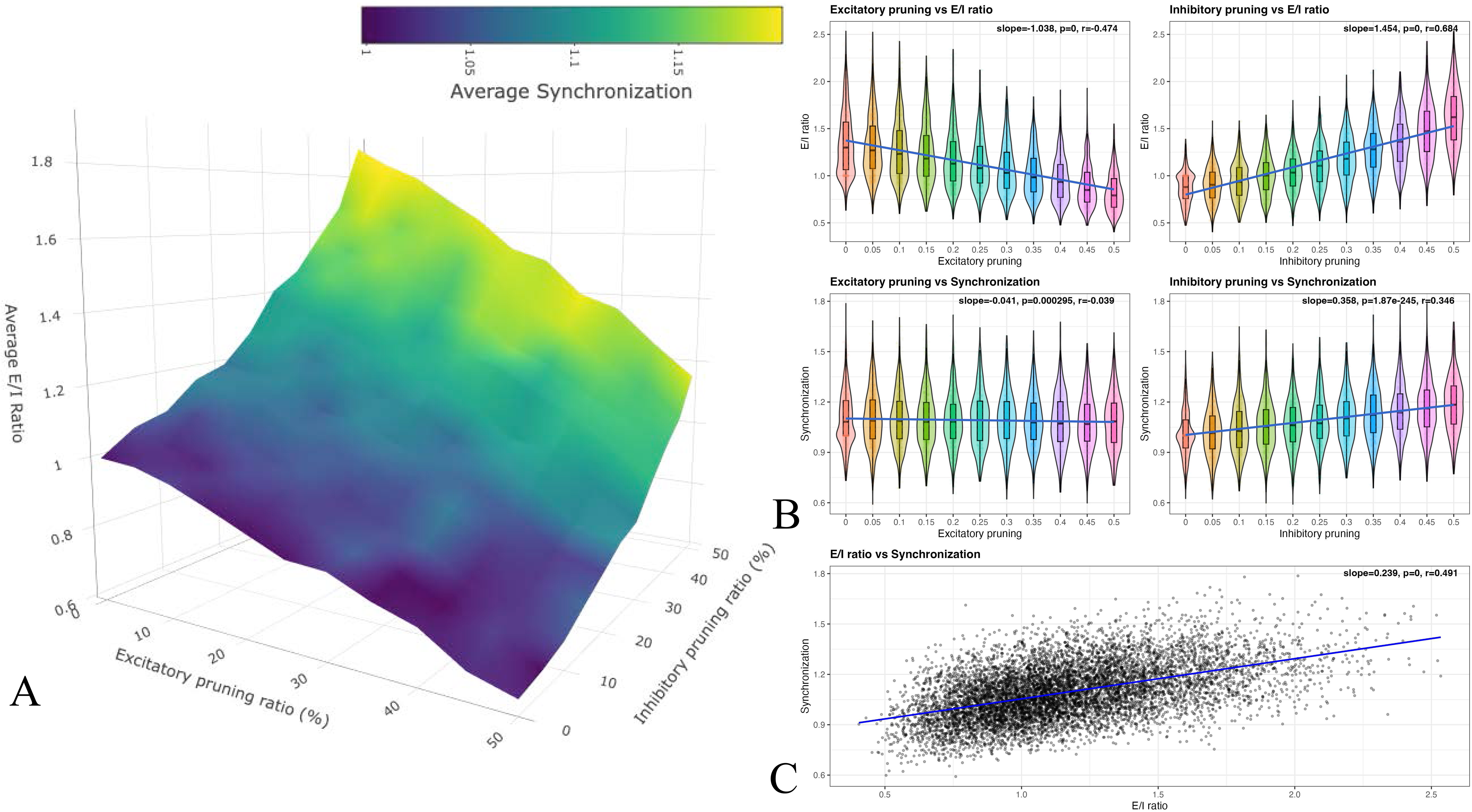
Neural circuit modeling of synaptic pruning. (A) Simulation results of excitatory pruning ratio (0-0.5), inhibitory pruning ratio (0-0.5), E/I ratio, and synchronization. X-axis: Excitatory pruning ratio; Y-axis: Inhibitory pruning ratio; Z-axis: Average E/I ratio. Colors indicate average synchronization levels from dark purple (close to 1, indicating no change in synchrony) to yellow (indicating higher synchrony). (B) The distribution of E/I ratio and synchronization values across different pruning levels. Each violin represents the distribution of values at a given pruning ratio (0–0.5 in 0.05 increments). Embedded boxplots display median and interquartile ranges. Overlaid dots represent individual simulation outcomes. Regression results (slope, p-value, and Pearson’s r) are shown in the upper right of each panel, indicating that inhibitory pruning has a stronger positive correlation with both the E/I ratio and synchronization. In contrast, excitatory pruning shows weaker correlations. (C) The E/I ratio and synchronization relationship across all simulation conditions. A positive correlation is observed, suggesting that changes in E/I balance mediate alterations in network synchrony. Regression statistics are displayed in the corner.

Our findings indicate that pruning excitatory or inhibitory synapses alters the E/I ratio, but with differential effects. Inhibitory synaptic pruning significantly elevated the E/I ratio, while excitatory pruning reduced it. These bidirectional trends were consistent across all simulations. In experiments where excitatory or inhibitory pruning was varied independently (**Fig. 4A**). Correlation analysis revealed a negative association between excitatory pruning and the E/I ratio (*r* = −0.474) and a stronger positive correlation between inhibitory pruning and the E/I ratio (*r* = 0.684) (Fig. 4B). Linear regression further confirmed this differential sensitivity, with excitatory pruning reducing the E/I ratio (β = −1.038). In contrast, inhibitory pruning increased more substantially (β *=* 1.454), indicating that E/I balance is more sensitive to changes in inhibitory connectivity (**Fig. 4B**).

Network synchronization exhibited an even more pronounced differential sensitivity to pruning type. Inhibitory pruning positively correlated with event synchronization (*r =* 0.346). In contrast, excitatory pruning showed only a negligible negative correlation (*r =* −0.039) (**Fig. 4B**). Linear regression analysis further confirmed this asymmetric sensitivity, with inhibitory pruning exerting a substantially more substantial effect on synchronization (β *=* 0.358) compared to excitatory pruning (β *=* −0.041) (**Fig. 4B**). Additionally, the E/I ratio itself was positively correlated with synchronization (*r =* 0.491), suggesting that the influence of synaptic pruning on network synchronization may be indirectly mediated through its effects on E/I balance (**Fig. 4C**).

These simulations demonstrate that while both types of synaptic pruning affect network properties, inhibitory pruning exhibits a heightened potency in disrupting both E/I balance and synchronization dynamics. This differential sensitivity to inhibitory pruning may represent a key mechanism by which synapse loss in neurodegenerative diseases may trigger the hypersynchrony characteristic of network hyperexcitability and epileptiform activity.

## 4. Discussion

While Alzheimer’s disease (AD) and epilepsy (EP) share a complex bidirectional relationship, the molecular mechanisms underlying their comorbidity remain poorly understood. Our genome-wide transcriptional analysis of multiple animal models and patient cohorts revealed several key molecular insights into this relationship, particularly regarding shared neuroinflammatory pathways and synaptic pruning mechanisms.

### Conserved Neuroinflammation Pathways in Epilepsy and Alzheimer’s Disease

Our genome-wide analysis proves that neuroinflammatory pathways form a common molecular substrate between AD and EP (22). We identified remarkable preservation of the immune-related turquoise module across disease models and human patient cohorts **(Fig. 1-2**). Functional enrichment analysis of the preserved module revealed significant enrichment in neuroinflammatory processes, such as “inflammatory response”, “cytokine production”, “blood vessel morphogenesis”, “glial cell activation”, and “complement and coagulation cascades” (**Table S2**).

Neuroinflammation has been extensively documented in both diseases and serves a complex dual role in disease progression (22,44,45). While activated microglia facilitate Aβ plaque clearance and reactive astrocytes provide metabolic support and ionic homeostasis that may attenuate neural damage and seizures, chronic activation or dysfunction of these glial populations can promote disease progression through persistent neuroinflammation (37,46,47). Specifically, activated microglia and astrocytes secrete proinflammatory cytokines and chemokines, including IL-1β, TNF-α, and IL-6, which can enhance neuronal hyperexcitability, increase seizure susceptibility, exacerbate Aβ and tau aggregation, and accelerate neurodegeneration and cognitive decline (48). Additional inflammatory components, including complement cascade activation and blood-brain barrier (BBB) dysfunction, further link these pathologies. Complement activation contributes to synaptic loss in AD and is elevated in TLE patient tissue. BBB disruption promotes both seizure susceptibility and neuroinflammation (49–52).

Our results supported previous findings that identified neuroinflammation as a key mechanistic link between AD and EP. AD-associated inflammation enhances seizure susceptibility, and EP-induced inflammation may accelerate Aβ and tau aggregation as well as cognitive decline.

### Synaptic Pruning as Core Signatures in this Comorbidity

Our analysis of the conserved immune module revealed a central synaptic pruning pathway involving *TYROBP*, *TREM2*, and *C1Q* complement genes (**Fig. 1**). These genes showed consistent co-dysregulation in microglia and demonstrated diagnostic value across both conditions (**Figs. 2-3**). The expression profile notably differentiated between activated and alternative metabolic state microglial populations (**Fig. 3I**). Collectively, these findings suggest that this microglia-mediated synaptic pruning may represent a core mechanistic pathway underlying the comorbidity of AD and EP.

This pathway is well-documented in AD pathology, with *TYROBP* and *TREM2* established as risk factors due to their effects on microglial function (53–55). Under physiological conditions, microglia maintain synaptic homeostasis through complement-mediated pruning, where *C1Q* tags unnecessary synapses for elimination (56). *TREM2* and *TYROBP* recognize these tags and initiate phagocytic clearance, which is critical for neural circuit maintenance (57). However, in AD pathology, these receptors trigger microglia’s transition to a disease-associated state (DAM), characterized by enhanced phagocytic activity and elevated expression of *TYROBP*, *TREM2*, and *C1Q* (57). While *TYROBP*/*TREM2* deficiency impairs plaque clearance, pathway overactivation can also promote neurodegeneration through excessive synaptic pruning (58–60).

The impact of synaptic pruning on AD and EP progression can operate through two primary mechanisms: synaptic transmission and plasticity. *C1Q* inhibition in aged models can restore both long-term potentiation (LTP) and depression (LTD), while increased *C1Q* expression impairs LTP in non-aged models (58–61). This is particularly relevant as impaired synaptic plasticity characterizes cognitive decline in AD and may contribute to epileptogenic circuit formation in EP. Elevated *C1Q* expression leads to decreased synaptic transmission in the CA1 region, affecting both excitatory and inhibitory neurons (67–69), which can create a harmful feedback loop where synaptic dysfunction promotes disease progression in both conditions (62,63).

Notably, interventions targeting the synaptic pruning pathway, whether through the knockdown of regulatory genes like *TYROBP*/*TREM2* or direct inhibition of *C1Q* accumulation, have successfully restored both electrophysiological and cognitive functions in AD models. This restoration occurred even in the presence of significant Aβ and tau pathology (57–60). Our results establish a regulatory network that is consistent with previous observations and further extends its role in the comorbidity of AD and EP (**Fig. 1B**). Interestingly, microglial clusters 1 and 6 showed downregulation of consensus signatures including *TYROBP*, *TREM2*, and *C1Q* genes (**Fig. 3I**). These clusters have been previously reported to correlate positively with amyloid-beta and tau tangle pathology, while correlating negatively with the slope of cognitive decline (40). Conversely, clusters 4, 9, and 10 upregulated these genes (*TYROBP*, *TREM2*, *C1Q*) and were enriched for genes that generally correlated negatively with amyloid-beta and tau tangle pathology, while correlating positively with the slope of cognitive decline (**Fig. 3I**) (40). These results may reflect the complex and stage-dependent role of microglia in the pathogenesis of AD. While microglia are designed to reduce the burden of AD pathology, their excessive activation may paradoxically contribute to neurodegeneration and cognitive decline.

While this pathway has been extensively studied in AD, its role in AD-EP comorbidity remains unexplored. To address this gap and understand how synaptic pruning affects global neural network dynamics in AD-EP comorbidity (64), we conducted an in silico experiment.

### Asymmetric Effects of Synaptic Pruning Promote Network Synchronization

Computational modeling has improved our understanding of AD and EP pathologies. It can predict quantitative mechanisms behind them, with models ranging from molecular-level ion dynamics simulations to large-scale network models (65,66). We developed a neural circuit model building upon Liu’s biologically plausible EP modeling framework (42). This model allowed us to quantify how pruning of excitatory and inhibitory synapses affects E/I balance (E/I ratio) and the temporal coordination of neural firing (synchronization).

Our model reveals a differential sensitivity of neural networks to synaptic pruning, with E/I balance being substantially more responsive to inhibitory synapse loss than excitatory synapse elimination (**Fig. 4**). Excitatory synapse elimination decreased the E/I ratio but had negligible effects on network synchronization. In contrast, inhibitory synapse pruning resulted in a significant increase in the E/I ratio and network synchronization (**Fig. 4**). This heightened sensitivity to inhibitory pruning can be explained by the architectural role of inhibitory interneurons within neural circuits. These neurons often establish synaptic connections with multiple excitatory neurons and act as crucial circuit coordination and synchronization regulators. Therefore, when inhibitory control mechanisms are impaired, dysregulation cascades throughout more widespread circuits (67). Consequently, a reduction in inhibitory synapses produces a disproportionately more significant perturbation in neural networks than an equivalent reduction in excitatory synapses. This result implies a plausible mechanistic pathway whereby neuronal or synapse loss in neurodegenerative diseases disrupts E/I balance, enhancing sensitivity to inhibitory synapse loss and thereby promoting EP susceptibility. (**Fig. 4**). This mechanism may extend beyond AD, as EP is a well-documented comorbidity of multiple neurodegenerative conditions, including Huntington’s, Parkinson’s disease, and multiple sclerosis (68). Notably, in AD, synaptic loss often precedes cognitive decline and EP, suggesting it may be an initiating event rather than a consequence (1).

Our results emphasize the critical role of inhibitory synaptic pruning in promoting network synchronization due to the neural network’s heightened sensitivity to inhibitory synapse loss. At the molecular level, *C1Q* functions as an “eat me” signal, tagging synapses for microglial elimination (64). While this tagging process is not synapse-type specific, interestingly, several recent studies have demonstrated that microglia may preferentially prune inhibitory synapses (64,69). Dejanovic et al. demonstrated that the *TREM2-C1Q* microglial synaptic pruning pathway engulfs inhibitory synapses more frequently than excitatory synapses in AD models. At the same time, Fan et al. observed similar preferential pruning of inhibitory synapses during recurrent seizures that exacerbate EP model pathology (69,70). Studies have shown that *C1Q* blockade reduces both the frequency and severity of seizures in traumatic brain injury models (71). However, it is also important to note that *C1Q* knockout also affects synaptic elimination during development, enhancing connectivity and seizure activity (72,73). Recently, Aloi et al. reported that overactivation of microglia reduced Aβ pathology and caused hyperexcitability and seizures via synaptic pruning (74).

These findings collectively suggest that the network’s differential sensitivity to inhibitory synapse loss and the apparent preferential pruning of inhibitory synapses by microglia may represent a critical mechanism underlying AD-EP comorbidity. Managing microglial *C1Q*-mediated synaptic pruning during AD pathology and aging warrants further investigation, particularly regarding regulating this potential preferential pruning of inhibitory synapses.

## Conclusion

Our genome-wide transcriptional analysis of AD and EP animal models and human cohorts revealed a consensus molecular signature converging on microglial synaptic pruning, with a highly preserved immune module centered on the *TYROBP-TREM2-C1Q* complement pathway consistently upregulated across species and conditions. To support community validation, we developed a web application (https://github.com/HuihongLi/OverlapModule) enabling researchers to test our module overlap preservation with theirs. Neural circuit modeling demonstrated that inhibitory synapse loss through complement-mediated pruning exerts disproportionately powerful effects on E/I balance and network synchronization compared to excitatory loss, creating a pathological loop of hyperexcitability and hypersynchrony that mechanistically links neuroinflammation to network dysfunction in AD-EP comorbidity. These findings provide a unifying framework explaining the bidirectional relationship between AD and EP, suggesting that targeting microglial-complement signaling may offer disease-modifying benefits.

## Abbreviations

EP: Epilepsy
AD: Alzheimer’s disease
E/I: Excitation/Inhibition
TLE: Temporal lobe epilepsy
Aβ: Amyloid-β
PO: Pilocarpine-induced
KA: Kainic acid-induced
GCNs: Gene co-expression networks
WT: Wild-type
PCA: Principal Component Analysis
BH: Benjamini-Hochberg
GO: Gene Ontology
TOM: Topological overlap matrices
MEs: Module eigengenes
MM: Module membership
GS: Gene-genotype significance
PPI: Protein-protein interaction
FDR: False discovery rate
AUC: Area Under the Receiver Operating Characteristic Curve
ROC: Receiver Operating Characteristic
CBE: Cerebellum
DLPFC: Dorsolateral prefrontal cortex
STG: Superior temporal gyrus
IFG: Inferior frontal gyrus
TCX: Temporal cortex
FP: Frontal pole
SVM: Support vector machine
ES: Event synchrony
BBB: Blood-brain barrier
DAM: Disease-associated microglia
LTP: Long-term potentiation
LTD: Long-term depression
ROSMAP: Rush Memory and Aging Project snRNA-seq Single nucleus RNA-sequencing

## Declarations

### Ethics approval and consent to participate

Not applicable.

### Consent for publication

Not applicable.

### Availability of data and materials

The datasets analyzed during the current study are available in the National Center for Biotechnology Information (NCBI), European Nucleotide Archive (ENA), and Synapse under the following accession numbers: PRJEB18790, GSE125957, GSE73878, GSE64398, GSE63808, E-MTAB-3123, GSE48350, GSE190453, syn51758062.1, GSE204702.

### Authors’ contribution

HL conceptualized the study, conducted the investigation, developed the methodology, performed formal analysis, and contributed to writing the original draft and reviewing/editing the manuscript. ZX, YT, RZ, YY, and BL contributed to the investigation, performed formal analysis, and participated in writing the original draft and reviewing/editing the manuscript. SC, ZD, and JL contributed to writing the original draft and reviewing/editing the manuscript. JW was involved in conceptualization and contributed to writing the original draft and reviewing/editing the manuscript. MC, XL, and YS contributed to the investigation and participated in reviewing/editing the manuscript. BH and NW conceptualized the study and contributed to writing the original draft and reviewing/editing the manuscript. XJ provided supervision, acquired funding, contributed to conceptualization, and participated in writing the original draft and reviewing/editing the manuscript. All authors read and approved of the final manuscript.

## Acknowledgments

The results published here are in whole or in part based on data obtained from the **AD Knowledge Portal.**

