## Supplementary material for "Genome-wide consensus transcriptional signatures identify synaptic pruning linking Alzheimer’s disease and epilepsy": Suppl Fig

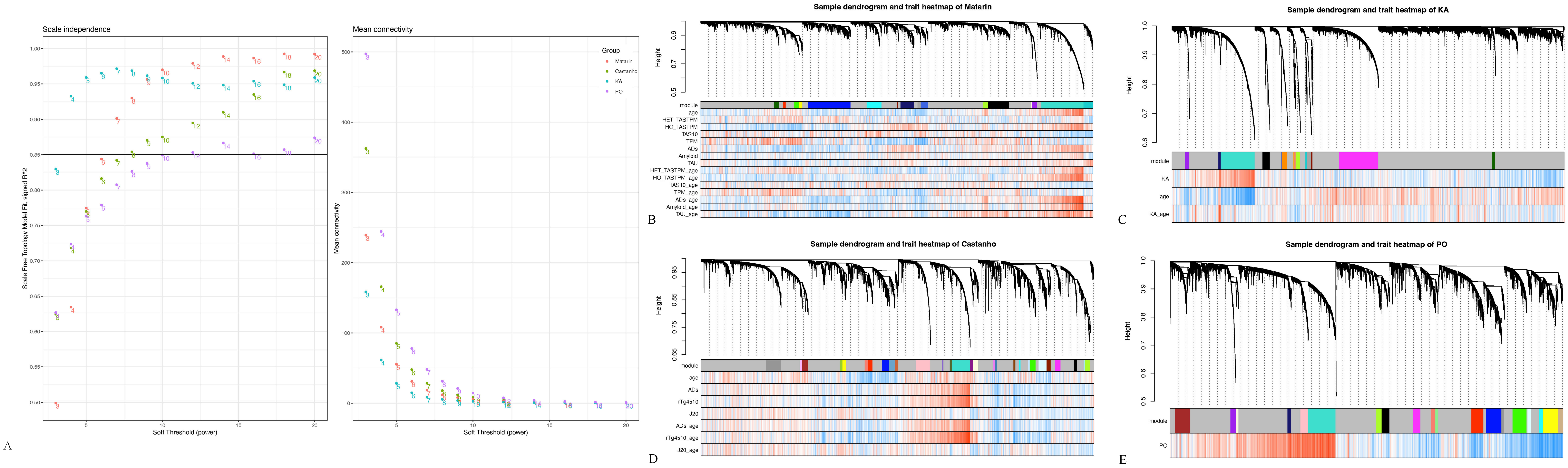
Scale independence
Mean connectivity
Group
Matarin
Castanho
KA
PO
Scale Free Topology Model Fit, signed R^2
Mean connectivity
Soft Threshold (power)
A
Sample dendrogram and trait heatmap of Matarin
Height
module
age
HET_TASTPM
HO_TASTPM
TAS10
TPM
ADs
Amyloid
TAU
HET_TASTPM_age
HO_TASTPM_age
TAS10_age
TPM_age
ADs_age
Amyloid_age
TAU_age
B
Sample dendrogram and trait heatmap of KA
module
KA
age
KA_age
C
Sample dendrogram and trait heatmap of Castanho
module
age
ADs
rTg4510
J20
ADs_age
rTg4510_age
J20_age
D
Sample dendrogram and trait heatmap of PO
module
PO
E

Figure S1

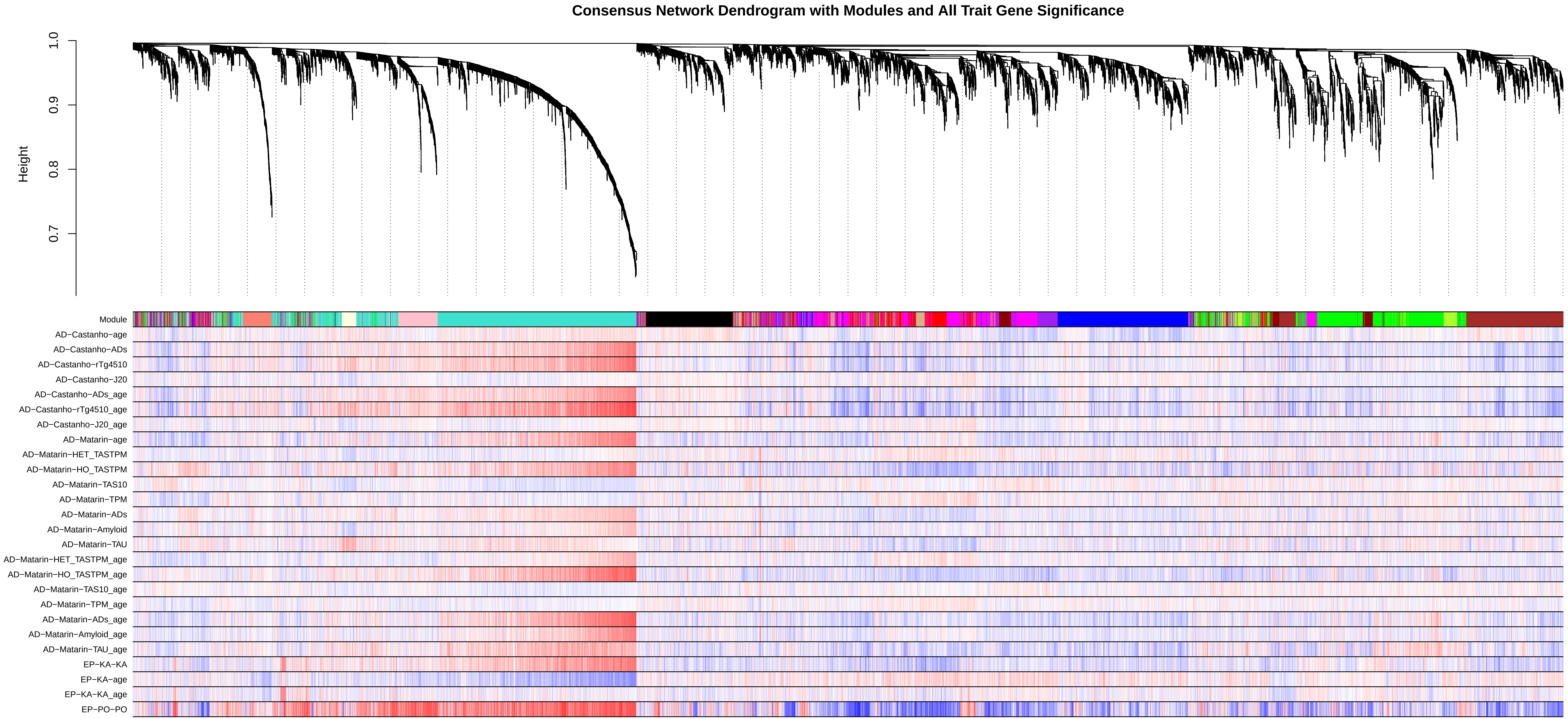
Consensus Network Dendrogram with Modules and All Trait Gene Significance
Height
1.0
0.9
0.8
0.7
Module
AD−Castanho−age
AD−Castanho−ADs
AD−Castanho−rTg4510
AD−Castanho−J20
AD−Castanho−ADs_age
AD−Castanho−rTg4510_age
AD−Castanho−J20_age
AD−Matarin−age
AD−Matarin−HET_TASTPM
AD−Matarin−HO_TASTPM
AD−Matarin−TAS10
AD−Matarin−TPM
AD−Matarin−ADs
AD−Matarin−Amyloid
AD−Matarin−TAU
AD−Matarin−HET_TASTPM_age
AD−Matarin−HO_TASTPM_age
AD−Matarin−TAS10_age
AD−Matarin−TPM_age
AD−Matarin−ADs_age
AD−Matarin−Amyloid_age
AD−Matarin−TAU_age
EP−KA−KA
EP−KA−age
EP−KA−KA_age
EP−PO−PO

Figure S2

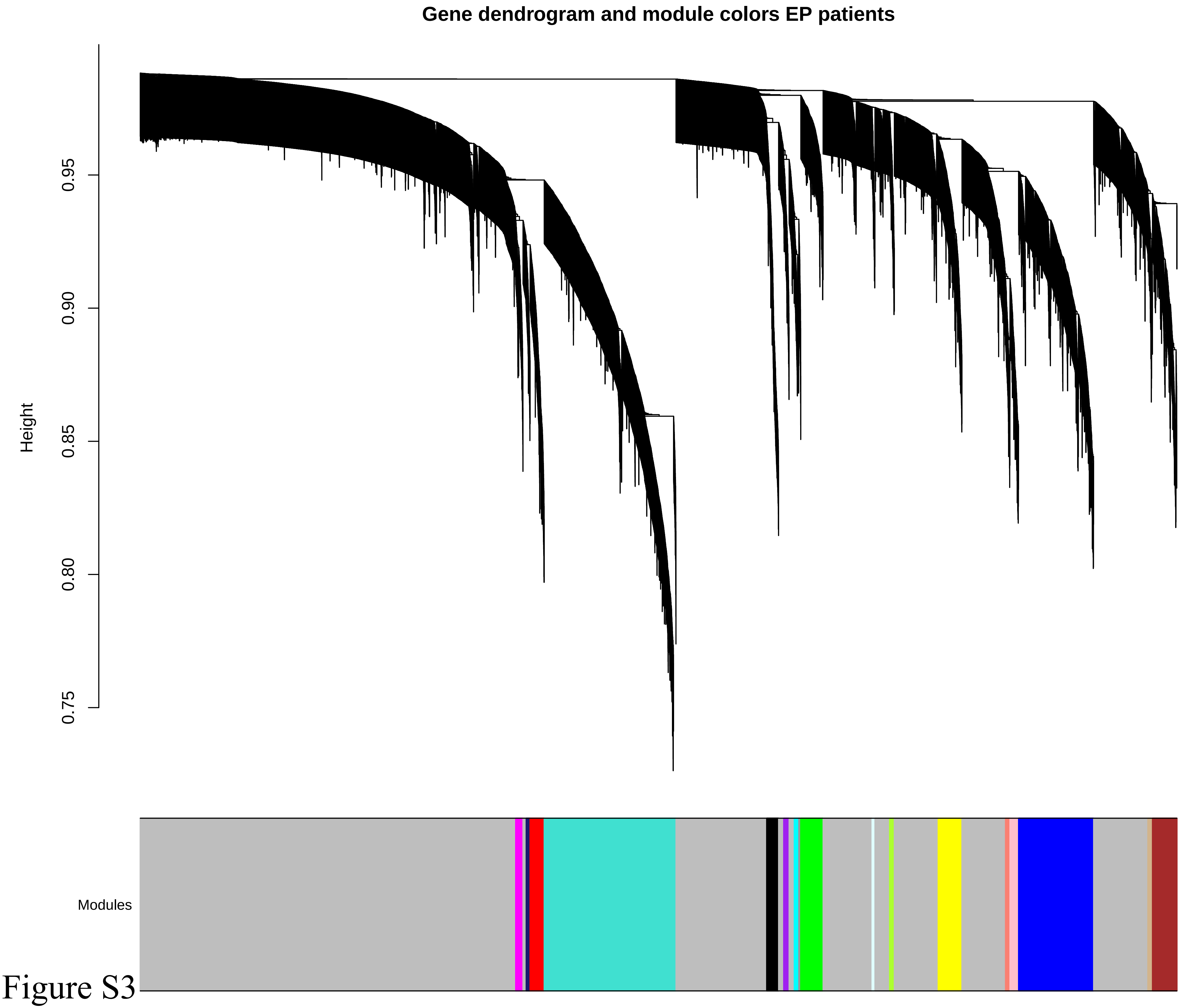
Gene dendrogram and module colors EP patients
Height
0.75
0.80
0.85
0.90
0.95
Modules

Figure S3

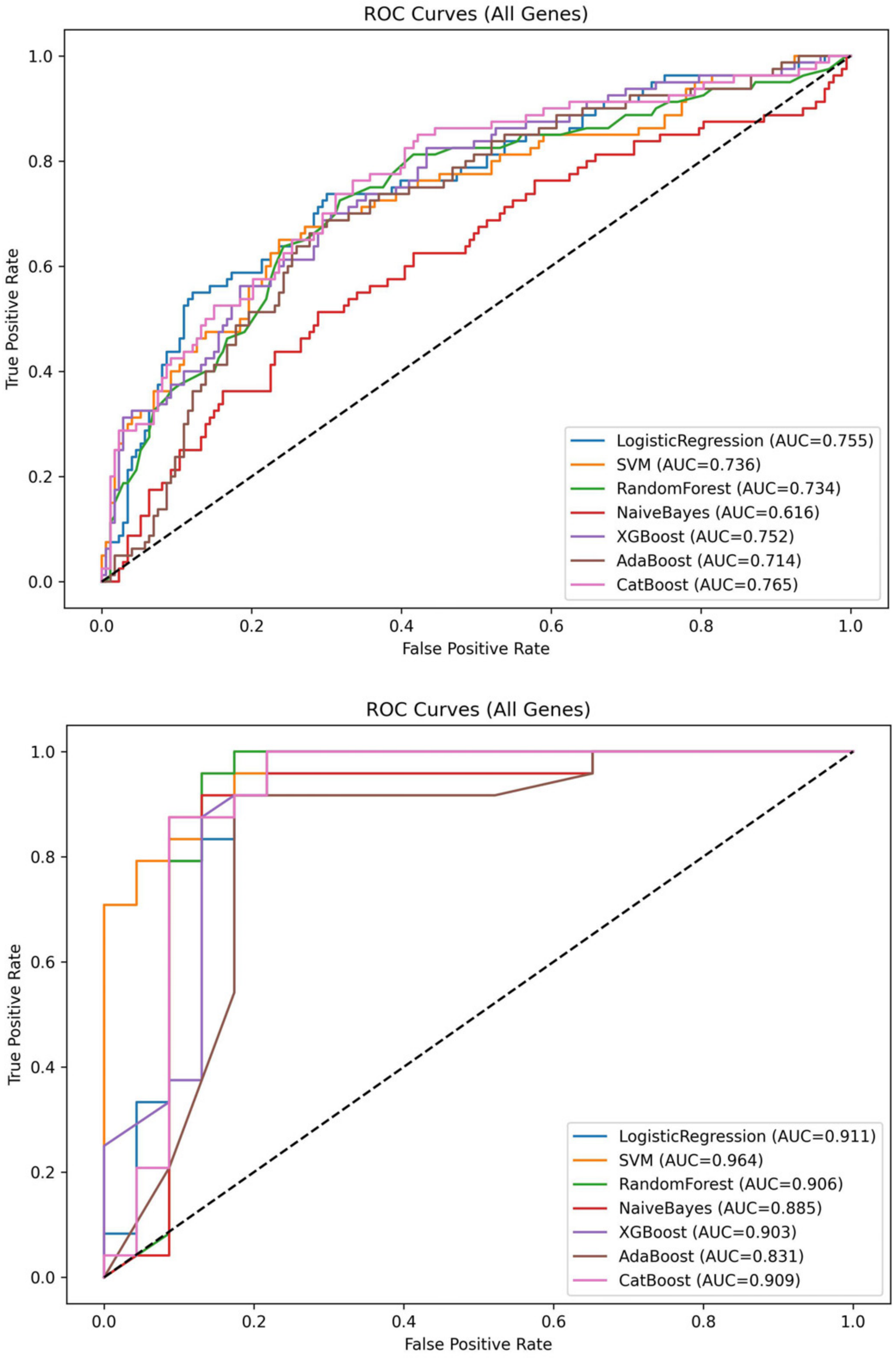
ROC Curves (All Genes)
True Positive Rate
False Positive Rate
LogisticRegression (AUC=0.755)
SVM (AUC=0.736)
RandomForest (AUC=0.734)
NaiveBayes (AUC=0.616)
XGBoost (AUC=0.752)
AdaBoost (AUC=0.714)
CatBoost (AUC=0.765)
ROC Curves (All Genes)
True Positive Rate
False Positive Rate
LogisticRegression (AUC=0.911)
SVM (AUC=0.964)
RandomForest (AUC=0.906)
NaiveBayes (AUC=0.885)
XGBoost (AUC=0.903)
AdaBoost (AUC=0.831)
CatBoost (AUC=0.909)

A

B

Figure S4

**Legends of supplementary files**

**Text S1: Supplementary Methods**

**Text S1: Model Details**

**Table S1: Summary of Quality Control and Mapping Statistics**

**Table S2: Statistics of Module Preservation and Functional Enrichment**

**Table S3. Module Gene List**

**Figure S1. Construction of co-expression modules in model datasets.**

(A) Scale independence and mean connectivity plots for different soft-thresholding powers across all four datasets (EP-PO, EP-KA, AD-Matarin, AD-Castanho). A power was selected as the minimum threshold that achieved scale-free topology ($R^2 \geq 0.85$) across all datasets. (B-E) Sample dendrograms and trait heatmaps showing hierarchical clustering of genes and resulting modules for (B) AD-Matarin, (C) EP-KA, (D) AD-Castanho, and (E) EP-PO datasets. Upper dendrograms show gene clustering, while lower heatmaps display module-trait relationships, with red indicating positive correlations and blue indicating negative correlations with disease status.

**Figure S2. Dendrogram of the consensus network**

The consensus network's hierarchical clustering dendrogram was constructed by integrating topological overlap matrices from all four datasets (EP-PO, EP-KA, AD-

Matarin, AD-Castanho). The color band beneath shows consensus module assignments, while the heatmap displays module-trait associations across multiple models, disease conditions, and age variables. This consensus approach identified robust co-expression patterns conserved across AD and EP models.

**Figure S3. Construction of co-expression module in EP patient datasets.**

Dendrogram of gene modules identified in the EP patient dataset using hierarchical clustering and dynamic tree cutting based on TOM. (parameters: Soft-thresholding power of 7 ensuring a scale-free topology ($R^2 \geq 0.85$), deepSplit = 1, minClusterSize = 27).

**Figure S4. ROC curves of diagnostic models using shared hub gene expression.**

ROC curves for classifiers trained on (A) the AD patient dataset (GSE48350) and (B) the EP patient dataset (E-MTAB-3123). Multiple machine learning classifiers, including Logistic Regression, Support Vector Machine (SVM), Random Forest, Naive Bayes, XGBoost, AdaBoost, and CatBoost, were evaluated using 5-fold stratified cross-validation. Models were trained using standardized expression values of all consensus hub genes.
