## Supplementary material for "Genome-wide consensus transcriptional signatures identify synaptic pruning linking Alzheimer’s disease and epilepsy": Text S1

**Data Collection**

We conducted a comprehensive search to identify relevant transcriptomic datasets from animal models and patient cohorts. The selection process involved screening based on sample size adequacy (n > 40) and research design to ensure statistical robustness and minimize bias. Our final selection comprised several large-scale, methodologically sound datasets from EP and AD research that focused on genome-wide expression patterns in well-characterized disease models and appropriate controls. This resulted in four large-scale animal model datasets, two EP patient cohorts, and one extensive AD patient cohort.

**Quality control and preprocessing**

All raw RNA sequences underwent quality control and trimming to obtain clean sequences, using FastQC (0.12.1) and Fastp (0.22.0) with stringent parameters to ensure high data quality (1). Ribodetector (0.2.7) was used to remove potential rRNA contamination (2). Trimmed sequences were then aligned to the GRCm39.111 reference mouse genome using HISAT2 (2.2.1) and followed by quantification using featureCounts (2.0.6) (3,4). Detailed information on mapping metrics is provided in **Table S1**. Principal Component Analysis (PCA) and hierarchical clustering were performed to identify potential outlier samples. All datasets underwent appropriate normalization: RNAseq data were processed with variance-stabilizing transformation using DESeq2 (1.46.0), while microarray data were subjected to log2 transformation and variance-stabilizing normalization using the "limma" (3.62.1) package (5,6). Batch effects were removed using limma:removeBatchEffect(). We assessed dataset comparability by calculating general network properties such as average gene expression and overall network connectivity, with results indicating satisfactory comparability across all datasets.

**Cell Composition Deconvolution**

In silico deconvolution can estimate the cellular composition of a tissue sample from its gene expression profile, with CIBERSORT particularly suited for cellular composition deconvolution of brain transcriptomes (7,8). Cell composition deconvolution was performed using CIBERSORT (1.04**)** via the BrainDeconvShiny platform, with cell-type signatures specific to *Mus musculus* from immunopurified mouse brain tissue (7,8). Associations between identified cell types and modules were determined by calculating Pearson correlation coefficients.

**Machine Learning**

To evaluate the diagnostic potential of consensus hub signatures, we conducted binary classification (disease vs. control) across datasets using these signatures as predictive features. Prior to analysis, gene expression values underwent standardization to ensure comparable scaling across genes. We implemented classification models in Python 3.8.20 using scikit-learn, XGBoost, and CatBoost libraries. Seven supervised learning algorithms were evaluated: Logistic Regression (with increased maximum iterations of 1000), Support Vector Machine with RBF kernel, Random Forest, Naive Bayes, XGBoost, AdaBoost, and CatBoost. To ensure robust performance estimation and prevent overfitting, we employed 5-fold stratified cross-validation, with each fold maintaining proportional representation of disease and control samples. Performance was assessed through multiple metrics, including Accuracy, Precision, Recall, F1 score, and Area Under the Receiver Operating Characteristic Curve (AUC). The AUC was calculated by pooling predicted probabilities from all cross-validation splits to generate Receiver Operating Characteristic (ROC) curves.
