## Supplementary material for "Genome-wide consensus transcriptional signatures identify synaptic pruning linking Alzheimer’s disease and epilepsy": Text S2

**Computational modelling**

To explore the contribution of the identified key pathways to pathological changes in AD and EP at the network level, we implemented a spiking neural network using the Izhikevich neuron model to simulate hippocampal CA3 circuit dynamics with simulation spans of 1000ms, employing the Brian2 package ^1^. The network consists of 1000 Izhikevich neurons (800 excitatory and 200 inhibitory), with neural dynamics governed by ^2^:

$$\frac{dv}{dt}=0.04v^{2}+5v+140-u+I$$

$$\frac{du}{dt}=a\left( bv-u \right)$$

where v is the membrane potential, u is the recovery variable. When v≥30 mV, the following reset conditions are applied: $v=c$; $u = u + d$. Parameters a, b, c and d are set to standard regular spiking for excitatory neurons and fast-spiking for inhibitory neurons following the Izhikevich framework ^2^. I is external input incorporating both suprathreshold Poisson noise at 1 Hz and presynaptic inputs ^3^.

Following Liu’s EP network modeling, synaptic connectivity is sparse, with a 0.1 probability of random connections between neurons, and interactions are modeled using conductance-based synapses to allow for voltage-dependent temporal and spatial summation ^3^. Synaptic interactions simulate AMPA (excitatory) and GABA-a (inhibitory) currents for four connection types: excitatory-excitatory (E→E), excitatory-inhibitory (E→I), inhibitory-excitatory (I→E), and inhibitory-inhibitory (I→I) ^2^. Synaptic currents are calculated as ^3^:

$$I_{j}=wS_{i\to j}\left( t \right)\left( E_{i}-v_{j} \right)$$

we introduce w to leverage the synaptic weight, while $S_{i\to j}\left( t \right)$) is the synaptic conductance evolving as $\frac{dS_{i\to j}\left( t \right)}{dt}=-\frac{S_{i\to j}\left( t \right)}{\tau\left( i \right)}$ and $E_{i}$ is the synaptic reversal potential ^3^. Synaptic conductance is updated upon presynaptic spikes by adding a peak conductance $S_{i\to j}\left( max \right)$, while we calibrate unitary synapse to induce EPSPs for E→E/E→I and IPSPs for I→E/I→I of 1.0, 2.5, −2.0, −1.0 mV with time constants:$\tau=\{3, 3, 5, 5\} \text{ms}$ and reversal potentials:$E=\{0, 0, -75, -75\} \text{mV}$ ^4-12^. Each of four types of synaptic weights (w) is drawn independently from a standard normal distribution, normalized to [0, 1], and set to 0 during synaptic pruning. We simulated synaptic pruning by selectively removing excitatory (E→E and E→I) or inhibitory (I→E and I→I) synapses, starting with those having the lowest weights based on the synaptic weight distribution (ranging from 0% to 50%, with 5% increments). Each pruning simulation was repeated at least 50 times.

The excitation/inhibition (E/I) ratio quantifies the balance between synaptic drives in the network, indicating firing rate stability, while event synchrony measures temporal coordination of firing patterns, implying network oscillation patterns. Both large E/I ratio and excessive synchrony are associated with EP and AD ^12^.

To analyze the network, we calculated the E/I ratio and event synchrony to measure the effects of synaptic pruning on firing rate stability and temporal coordination. The E/I ratio compares total excitatory currents (from E→E and E→I synapses) to the absolute magnitude of inhibitory currents (from I→E and I→I synapses), averaged over the simulation period T ^3^:$\text{ }$

$$E/IRatio=\frac{\sum_{i=1}^{N} I_{i}^{E}\left( t \right)}{\left| \sum_{i=1}^{N} I_{i}^{I}\left( t \right) \right|}$$

where:

${I_{i}}^{E}\left( t \right)$represents excitatory synaptic currents onto neuron i at time t

${I_{i}}^{I}\left( t \right)$ represents inhibitory synaptic currents onto neuron i at time t

N is the total number of neurons

$\bar{x}$represents the mean over time T

Event synchrony (ES) quantifies spike timing coordination between excitatory neuron pairs by calculating the fraction of their spikes occurring within 5-ms windows. For each excitatory neuron pair (i, j), synchronous events are counted (with exact coincident spikes weighted by 0.5 and near-coincident spikes within the time window weighted by 1.0), then normalized by the geometric mean of both neurons' total spike counts. During total simulation time T ^13^:

$$ES_{ij}=\frac{\sum_{s_{i}\in S_{i}} \sum_{s_{j}\in S_{j}} C\left( s_{i},s_{j} \right)}{\sqrt{N_{i}N_{j}}}$$

where:

Si is the set of all spike times from neuron i during the simulation.

Sj is the set of all spike times from neuron j during the simulation.

Ni and Nj are the total number of spikes from neuron i and j respectively over the entire simulation period.

C(si, sj) counts synchronized events for each pair of spikes:

$$C\left( s_{i},s_{j} \right)=\left\{ \begin{matrix} 1 & if 0<|s_{i}-s_{j}|<5ms \\ 0.5 & ifs_{i}=s_{j} \\ 0 & otherwise \end{matrix} \right.$$

The network-wide event synchrony is computed as ^13^:

$$\text{Network ES}=\frac{1}{P}\sum_{i,j} ES_{ij}$$

Where $P=\frac{Ne (Ne-1)}{2}$ is the total number of unique excitatory neuron pairs, Ne is the number of excitatory neurons.

Both E/I ratio and ES values are normalized to baseline unpruned values to highlight pruning-induced changes in network dynamics. After normalization, the E/I ratio and ES value fluctuate around 1. A value greater than 1 means a higher E/I ratio or ES value than after synaptic pruning, and a value less than 1 means a smaller E/I ratio or ES value after synaptic pruning. For statistical analysis, we conducted a correlation analysis by calculating Pearson correlation coefficients and linear regression analysis to quantify the effect sizes of excitatory and inhibitory pruning on E/I ratio and synchronization.
